# Heritable functional architecture in human visual cortex

**DOI:** 10.1101/2020.05.04.077545

**Authors:** Ivan Alvarez, Nonie J. Finlayson, Shwe Ei, Benjamin de Haas, John A. Greenwood, D. Samuel Schwarzkopf

## Abstract

How much of the functional organization of our visual system is inherited? Here we tested the heritability of retinotopic maps in human visual cortex using functional magnetic resonance imaging. We demonstrate that retinotopic organization shows a closer correspondence in monozygotic (MZ) compared to dizygotic (DZ) twin pairs, suggesting a partial genetic determination. Using population receptive field (pRF) analysis to examine the preferred spatial location and selectivity of these neuronal populations, we estimate a heritability around 30% for polar angle preferences and spatial selectivity, as quantified by pRF size, in extrastriate areas V2 and V3. Our findings are consistent with heritability in both the macroscopic arrangement of visual regions and stimulus tuning properties of visual cortex. This could constitute a neural substrate for variations in a range of perceptual effects, which themselves have been found to be at least partially genetically determined. These findings also add convergent evidence for the hypothesis that functional map topology is linked with cortical morphology.

**Highlights:**

- We analyzed retinotopic maps from monozygotic and dizygotic twin pairs
- Visual field maps in V1-V3 are more similar in monozygotic twins
- Heritability is greater in V1 and V3 for polar angle and population receptive field sizes
- Eccentricity maps show lesser degree of heritability
- Further evidence for link between cortical morphology and topology of retinotopic maps

**Graphical Abstract:** 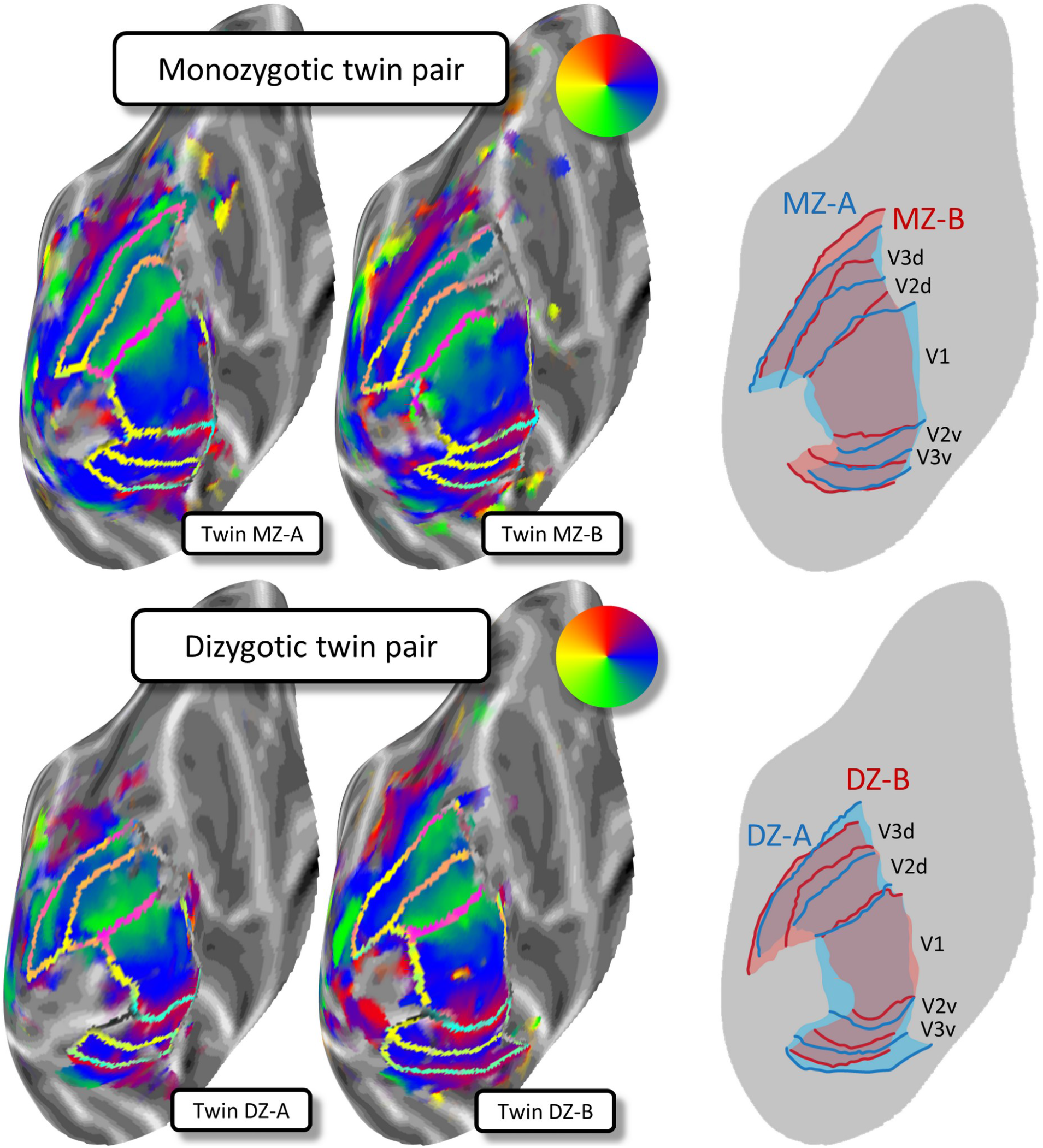

## 1. Introduction

Many aspects of visual perception show pronounced individual differences. These variations have further been shown to be at least partly genetically determined in processes including binocular rivalry (Miller et al., 2010), bistable perception (Shannon et al., 2011), eye movement patterns (Constantino et al., 2017; Kennedy et al., 2017), and even complex functions like face recognition (Wilmer et al., 2010; Zhu et al., 2010). This mirrors the heritability in the coarse structural and morphological features of the brain (Chen et al., 2011; Jansen et al., 2015), as well as the topology of functional networks including those in the visual cortex (Anderson et al., 2021). A genetic component has also been reported for aspects of cortical function thought to underlie visual processing. For instance, studies using magnetoencephalography have shown that the peak frequency of visually-induced gamma oscillations in early visual cortex is heritable (van Pelt et al., 2012). Such oscillations may derive from the local circuitry of neuronal populations (Pinotsis et al., 2013) that in turn relate to the macroscopic cortical morphology (Gregory et al., 2016; Schwarzkopf et al., 2012).

Perhaps the most striking property of the human visual system is its functional organization into retinotopic maps, where adjacent locations in the visual field map onto adjacent neuronal populations in visual cortex (Glickstein and Whitteridge, 1987). This organization further determines the borders of individual regions in human visual cortex (Engel et al., 1997; Sereno et al., 1995; Wandell et al., 2007). Both intrinsic genetic and extrinsic afferent processes govern how the cortex differentiates into different regions during development (O’Leary et al., 2007; Rakic et al., 2009). Retinotopic map organization also shows some correspondence with cortical morphology and microstructure (Benson et al., 2012; Sereno et al., 2013). Given these links, it follows that the organizational principles of visual cortex and its fine-grained functional properties may also be partially heritable. However, to date this remains untested.

Twin studies provide a unique opportunity to study environmental and genetic factors for individual differences in the architecture of the human visual system. The genetic proportion of observed variation between humans is defined as *heritability* (Falconer, 1965; Jansen et al., 2015), which is distinct from environmental influences, either shared or unique. Identical (MZ) twins share 100% of their genes, whereas non-identical (DZ) twins share 50% on average. Classical twin designs compare correlations between MZ and DZ twin pairs to isolate how much variance in a variable of interest can be attributed to genetic versus environmental components (Falconer, 1965).

Here, we set out to understand how much of the retinotopic organization of human visual cortex can be explained by genetic factors. We examined broad characteristics of the functional architecture of early visual cortex by conducting retinotopic mapping experiments with functional magnetic resonance imaging (fMRI) on both MZ and DZ twins. Specifically, each participant underwent three 8-minute runs of fMRI in which they viewed stimuli combining a rotating wedge with expanding or contracting rings (Alvarez et al., 2015; Stoll et al., 2020). We then used population receptive field (pRF) analysis (Dumoulin and Wandell, 2008; Moutsiana et al., 2016) to estimate the preferred visual field location and spatial selectivity of each voxel in visual cortex. We used an analysis approach with high statistical power that effectively treats each twin pair as an independent replication. We provide the first evidence that the fine-grained organization of retinotopic maps in early regions V1-V3 is more similar in MZ than DZ twin pairs, suggesting a genetic component. Specifically, we estimate the heritability of polar angle preferences and spatial selectivity in extrastriate areas V2 and V3 to be around 30%.

## 2. Materials and Methods

### 2.1 Participants

We collected data from 36 pairs of same-sex twins. All participants were healthy and had normal or corrected-to-normal visual acuity. Participants were financially compensated for their time and travel costs. We obtained written informed consent from all participants and all procedures were approved by the University College London Research Ethics Committee.

*Monozygotic (MZ)*. We recruited 22 pairs of monozygotic (MZ) twins (18 female, 4 male), mean age 25.1 years (18-47 years). Fifteen pairs were right-handed, one pair was left-handed, six pairs were mixed (one twin right-handed, one left-handed). fMRI data from three MZ pairs was excluded: one pair was excluded due to an incidental neurological finding in one twin, with two pairs excluded due to excessive head movement during scanning. All analyses presented here used the remaining 19 MZ pairs.

*Dizygotic (DZ)*. We recruited 14 pairs of dizygotic (DZ) twins (10 female, 4 male), mean age 25.0 years (18-40 years). Thirteen pairs were right-handed, and one pair was left-handed. All DZ twins are included in the analyses presented here.

#### 2.1.1. Demographics and questionnaire responses

Participants were also asked to complete a questionnaire to provide information about environmental factors in their upbringing, such as how often they shared school classes and friends, dressed alike as children, and how often they keep in contact with their twin (see Supplementary Information for the full questionnaire). Crucially, in addition to the self-report of zygosity from the twins (Question 2), this questionnaire also included two questions previously validated to classify twin zygosity with 95% accuracy (Sarna et al., 1978):

> *During childhood, were you and your twin as alike as ‘two peas in a pod’ or were you of ordinary family likeness?* (Question 3)
>
> *Were you and your twin so similar in appearance at school age that people had difficulty in telling you apart?* (Question 4)

Self-report of twin zygosity originally led to 17 MZ and 19 DZ pairs of twins. The questionnaire results found conflicting categorizations, where the questionnaire-criteria zygosity for some twin pairs conflicted with self-reported zygosity. Five DZ pairs disagreed on both criteria (Questions 3 *and* 4), and three DZ and two MZ pairs disagreed on one criterion (Question 3 *or* 4). These 10 twin pairs were contacted and asked if they would take a genetic test through a third-party company (NorthGene, UK). Seven of the eight DZ pairs took the test, and of those, genetic testing found that the five who disagreed on both criteria were (probable) MZ twins. The two (of three) DZ pairs who disagreed on one criterion were both DZ pairs. None of the MZ twin pairs took the genetic test. We consequently switched the categories for the DZ twins whose genetic testing indicated they were MZ twins, leaving us with 22 MZ and 14 DZ twin pairs. This reclassification was done independently, prior to the main data analysis.

### 2.2. Functional MRI experiment

#### 2.2.1. Parameters

Imaging data were collected on a Siemens Avanto 1.5T MRI scanner located at the Birkbeck-UCL Centre for NeuroImaging, using a 32-channel head coil with the two coils in the middle of the top half of the head coil restricting vision removed, leaving 30 channels. Functional data were acquired with a T2*-weighted multiband 2D echo-planar sequence (2.3 mm isotropic voxels, 36 slices, FOV = 96×96 voxels, TR = 1s, TE = 55ms, acceleration factor = 4). Slices were oriented to maximize coverage of occipital cortex, generally approximately parallel to the calcarine sulcus. Each participant completed three functional runs mapping population receptive fields (pRF) with 490 volumes per run (including 10 dummy volumes), and two functional runs for localizing face and scene regions (not reported here). A high-resolution T1-weighted MPRAGE structural scan (voxels = 1mm isotropic, TR = 2730ms, TE = 3.57ms) was also obtained for each participant.

#### 2.2.2. Stimuli and task

Each scanning session lasted approximately 1 hour. Participants lay supine with the stimuli projected onto a screen (resolution: 1920 × 1080) at the back of the bore, via a mirror mounted on the head coil. The total viewing distance was 68cm.

We used a wedge and ring stimulus containing colorful images to map participants’ visual field locations. The wedge subtended a polar angle of 12° and rotated in 60 discrete steps (one per second). The maximal eccentricity of the ring was 8.5° and expanded/contracted over 36 logarithmic steps. Within each run, there were 6 cycles of wedge rotation and 10 cycles of ring expansion/contraction, interleaved with a 30 s fixation-only period after every quarter of the run. This stimulus rotated, expanded, and contracted around a central fixation dot. The order of rotation and expansion/contraction was the same in each run. The wedge and ring apertures contained previously described natural images (Moutsiana et al., 2016) or phase-scrambled versions thereof. Every 15 s the stimuli alternated between intact and phase-scrambled images. The sequence of individual images was pseudo-randomized. Participants were instructed to fixate at all times, and press a button if the fixation dot changed color from black to red (it could also change to a range of other colors) or if they saw a tartan pattern appear within the wedge and ring stimulus.

To analyze the behavioral data, for each participant and scanning run, we counted any time that the participant pressed the response button within 400 ms of an event (fixation color change or appearance of the tartan pattern) as a hit. Button presses outside those times were counted as false alarms. We then calculated the sensitivity, d’, of these response rates and averaged them across the three scanning runs. The results are shown in Supplementary Figure S1. There was no significant difference in behavioral sensitivity between twin groups (t(64)=-0.3078, p=0.7592), which suggests that both groups performed the task at comparable levels. Crucially, behavioral performance was also not significantly correlated between twins in each pair in either group (MZ: r=0.19,p=0.441; DZ: r=0.37,p=0.188). This means that our findings in the retinotopic mapping analysis are unlikely to be explained by more similar behavioral performance (e.g. through greater vigilance) in MZ than DZ twins.

#### 2.2.3. Pre-processing and pRF modelling

Functional MRI data were pre-processed using SPM12 (http://www.fil.ion.ucl.ac.uk/spm), MATLAB (MathWorks), and our custom SamSrf 5 toolbox for pRF mapping (https://doi.org/10.6084/m9.figshare.1344765). The first 10 volumes of each run were removed to allow the signal to reach equilibrium. Functional images were mean bias corrected, realigned and unwarped, and co-registered to the structural scan, all using default SPM12 parameters. FreeSurfer v5.3.0 (https://surfer.nmr.mgh.harvard.edu/fswiki) was used for automatic segmentation and reconstruction to create a 3D inflated model of the cortical surface from the structural scan. Functional data were projected onto the reconstructed cortical surface mesh, by sampling for each mesh vertex the time course from the nearest voxel midway between the white and grey matter surface. Linear trends were removed and time courses were *z*-normalized. The time courses of the three pRF mapping runs were averaged. Only vertices in the occipital lobe were included for further analyses, and all further analyses were performed in surface space.

Three parameters of a symmetrical, two-dimensional Gaussian pRF model were estimated for each voxel independently: *x*_*0*_, *y*_*o*_, and *σ*, where the first two denote the center coordinates of the pRF in the visual field and the third is the estimate of pRF size (standard deviation of the Gaussian). The model predicted the neural response at each time point of the fMRI time course from the overlap between the pRF model and a binary mask of the visual stimulus; the resulting time course was then convolved with a canonical hemodynamic response function. We then found the combination of pRF parameters whose time course best predicted the measured time course. Various descriptions of the data were then derived from these parameters, including: *polar angle, eccentricity*, and *R*^*2*^ (proportion variance explained).

We conducted pRF model fitting in two stages. First, a coarse fit using an extensive grid search was performed on data smoothed with a large Gaussian kernel on the spherical surface (FWHM=5 mm). The best fitting pRF parameters were determined as those producing the maximal Pearson correlation between the predicted and observed fMRI time course. Then we conducted a fine fit, using parameters identified by the coarse fit on a vertex by vertex basis to seed an optimization algorithm (Lagarias et al., 1998; Nelder and Mead, 1965) to minimize the sum of squared residuals between the predicted and observed time course. Only vertices whose goodness of fit on the coarse fit exceeded R^2^>0.05 were included in the fine fit. This stage used the unsmoothed functional data and also included a fourth amplitude parameter to estimate response strength. The final estimated parameter maps were then again smoothed on the spherical surface (FWHM=3 mm).

#### 2.2.4. Spatial normalization and manual delineation of visual regions

In order to compare retinotopic maps directly across different participants, we aligned all individual surfaces (and the final smoothed retinotopic maps) to the common space of the FreeSurfer *fsaverage* template. We calculated an average retinotopic map separately for each group, MZ and DZ, respectively. Then we averaged these two group maps together into one grand average map. This minimizes the undue influence the MZ group could have had on the average map due to its larger sample size. We then delineated visual regions V1, V2, and V3 based on reversals in the polar angle map and the extent of the activated portion of visual cortex along the anterior-posterior axis. Furthermore, we delineated the maps of all individual participants. This delineation was conducted blind with regard to the zygosity by presenting the maps of individual MZ and DZ in a shuffled order and hiding any identifying information from the analyst. A second analyst also delineated the regions (again blinded with regard to zygosity) using the retinotopic maps in native space without spatial normalization.

Data in regions V3A, V3B, and V4 were less consistent across participants, and particularly susceptible to variable signal-to-noise ratios between participants. Therefore, we did not analyze data from these regions further and restricted all of our analyses to V1-V3 only.

#### 2.2.5. Calculating regional overlap

To compare the extent of overlap in each visual regions for the twins in a pair we calculated an overlap score (Jaccard coefficient). This analysis requires that brains from individuals are first brought into register. We therefore used spatially-normalized surface maps for each participant, generated by cortical alignment to a common template brain. Specifically, for each twin pair, we determined the number of vertices in the reconstructed surface mesh that belonged to a given visual region in either twin. We further quantified the proportion of those vertices that overlapped between the two twins in the pair (area of overlap divided by the area of union). This index is therefore 1 when the two regions are exactly identical and 0 if the regions do not overlap. Any differences in the spatial relationship between a given region in two individuals will be captured in this range from 0-1, including the situation where the region for one twin is contained within that of the other. We then compared the region overlap scores between twin groups and visual regions.

#### 2.2.6. Calculating map similarity

Regional overlap is only a coarse measure of the topology of the cortical surface. Theoretically, it is possible that the shape of these regions could be identical in a twin pair but the retinotopic map organization within the regions could nevertheless be completely different. Regional overlap measures are also strongly dependent on the manual definition of regions, even if those definitions were performed blinded with regard to zygosity. To quantify map similarity, we calculated how pRFs in a given twin pair differed, taking all three pRF parameters into account. Specifically, for each vertex we calculated the Euclidean distance between the pRFs from each twin in the pair in a three-dimensional space defined by the x- and y-position, and the pRF size. We then averaged these distances across each visual region for each pair and took the inverse, converting distance into a similarity score. Finally, we compared these scores between twin groups and visual regions. Statistical inference was conducted using the logarithm of these scores to ensure linearity.

#### 2.2.7. Correlation analysis of pRF parameters

To analyze the similarity in individual pRF parameters separately, we then used the data extracted from each region to calculate the Spearman correlation of eccentricity and pRF size, and the circular correlation for polar angle, respectively, between twins in each pair. It is crucial to remove global trends before conducting such an analysis; otherwise any correlation between twins could simply be due to the pattern shared by all participants. For instance, the gradient of the eccentricity map will generally increase along the posterior-anterior axis. We therefore first removed the global trend from these data by computing the mean pattern of pRF parameters for the whole participant sample, and subtracting this mean pattern from the data of each individual. The pair-wise correlation was then carried out using the resulting residual parameters.

This analysis results in a set of correlation coefficients – for each pRF parameter there is one coefficient per twin pair in each group. Next, we calculated the average correlation across the twin pairs within each twin group (after Fisher’s z-transformation to linearize the correlation coefficients). We used bootstrapping to determine the 95% confidence intervals of these group averages by resampling the correlation coefficients from twin pairs 10,000 times with replacement and determining the 2.5^th^ and 97.5^th^ percentiles of the resulting distribution. This effectively determines the strength and reliability of the correlation at the group (i.e., between-subject) level. Heritability (H^2^) was calculated for each bootstrap iteration using Falconer’s formula (Falconer, 1965) given by *H*^*2*^ *= 2(M(r*_*MZ*_*) – M(r*_*DZ*_)), where *M(r*_*MZ*_) and *M(r*_*DZ*_) stand for the mean correlation coefficients in the MZ and DZ groups, respectively.

Finally, we conducted statistical inference on this using the bootstrapped data. To test if the heritability within a given visual region was significantly greater than zero, we quantified the proportion of bootstrap iterations where heritability was ≤0. To determine if heritability increased across the visual hierarchy, we fit a linear regression to the heritability values (regions dummy coded as 1, 2, and 3) for each bootstrap iteration. We then determined the significance of this regression by quantifying the proportion of iterations where the slope was ≤0.

#### 2.2.8. Power analysis

We conducted a simulation to estimate the statistical power of the correlation analysis (Supplementary Figure S2). To this end, we simulated data with the same number of spatial points as V3, the smallest visual region that we report (3409 vertices), and the same sample sizes for the MZ (n=19) and DZ (n=14) groups as in our study. These simulated data sets were drawn from a bivariate Gaussian distribution centered on zero and with a pre-specified correlation. We varied the difference in correlation between the MZ and DZ groups to simulate a range of true heritability values. Starting with r_MZ_=r_DZ_=0.5, which means zero heritability, we then gradually increased r_MZ_ by 0.001 and decreased r_DZ_ by the same amount, until we reached 0.53 (h^2^=12%). The correlation of 0.5 was chosen to be a reasonable mid-point of similarities. Because of how heritability is defined, only the difference between MZ and DZ intra-class correlations matters; thus, the same level of heritability can result from different baseline levels of correlation between DZ twins.

However, the baseline level of the correlation does have some impact on statistical power. To quantify the absolute minimal statistical power needed, we repeated this power simulation with a scenario with very low correlations. Specifically, we fixed r _DZ_=0 and then gradually increased r_MZ_ in steps of 0.001 until we reached 0.06. This again corresponds to (h^2^=12%) but statistical power was reduced relative to the other scenario.

We repeated these simulations 1,000 times. For each simulated heritability value, we then quantified the number of times that our bootstrapped heritability analysis detected a significant effect (p<0.05, Bonferroni corrected by 9 comparisons).

## 3. Results

### 3.1. Greater similarity in retinotopic map organization for MZ than DZ twins

We first consider the heritability of broad characteristics of the retinotopic maps in early visual areas. Figure 1 shows example retinotopic maps from one MZ twin pair (Figure 1A) and one DZ twin pair (Figure 1B) on the occipital lobe of the template brain. The general topology of the visual regions is more similar in the MZ pair than the DZ pair, which can be seen in the overlaid maps where borders between the visual areas are more closely aligned for the MZ (Figure 1C) than the DZ (Figure 1D) twins. Differences in this DZ pair are most evident in ventral occipital cortex (lower part of panel D), although there are also visible discrepancies for the dorsal V2/V3 border. For the MZ twins, differences are most pronounced in the dorsal regions, particularly towards the peripheral visual-field representation (top-right of panel C). Additional examples and maps for eccentricity and pRF size can be seen in the Supplementary Information.

**Figure 1.**
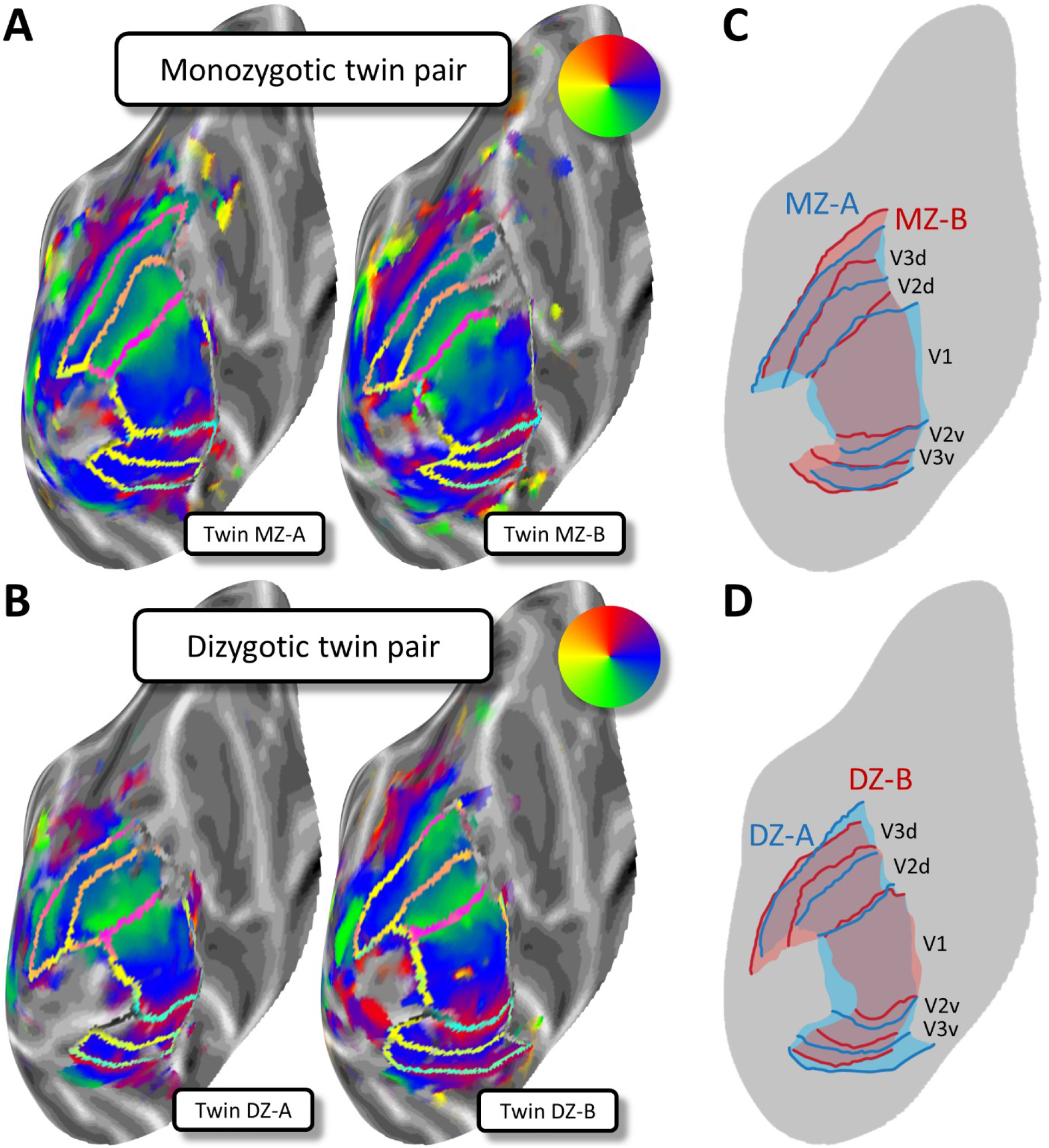
*A-B*. Retinotopic polar angle maps for an identical MZ (A) and a non-identical DZ twin pair (B). Polar angle maps are shown on an inflated model of the cortical surface for the left occipital lobe. Cortical surfaces were normalized by aligning them to a common template. Each plot shows data from one individual. Greyscale indicates the cortical curvature, with darker patches corresponding to sulci, and lighter patches corresponding to gyri. The pseudo-color code (see insets) denotes the preferred polar angle in the visual field for a voxel at a given cortical location, as derived from population receptive field (pRF) analysis. The transparent borders show transition boundaries for visual areas V1, V2 and V3. *C-D*. Schematic retinotopic maps with the borders for both twins in the MZ (C) and DZ (D) twin pairs overlaid in distinct colors to allow a direct comparison.

To quantify the topological similarity of these visual regions, we analyzed the overlap (Jaccard coefficient) of regions V1-V3, based on manual delineations of the retinotopic maps normalized to the template surface. To exclude experimenter bias, we carried out the delineation blinded to the zygosity of each participant. Figure 2A shows that, on average, the overlap across all three regions was consistently greater in MZ than DZ twins (two-way analysis of variance, main effect of zygosity: F(1,93)=18.29, p<0.0001). The amount of overlap also decreased from V1 to V3 (main effect of region: F(2,93)=151.22, p<0.0001). Variability in the topographical location of these visual regions is known to increase along the cortical visual hierarchy (Benson et al., 2012; Wang et al., 2015), and thus, the delineations of borders in extrastriate areas V2 and V3 are likely to be more variable than for region V1. The overlap of a particular region also depends on the borders with neighboring regions, making the results for the three regions partially dependent. Nevertheless, the difference between twin types did not differ significantly across the visual regions, with no significant interaction between the factors for visual region and zygosity (F(2,93)=0.22, p=0.8035).

**Figure 2.**
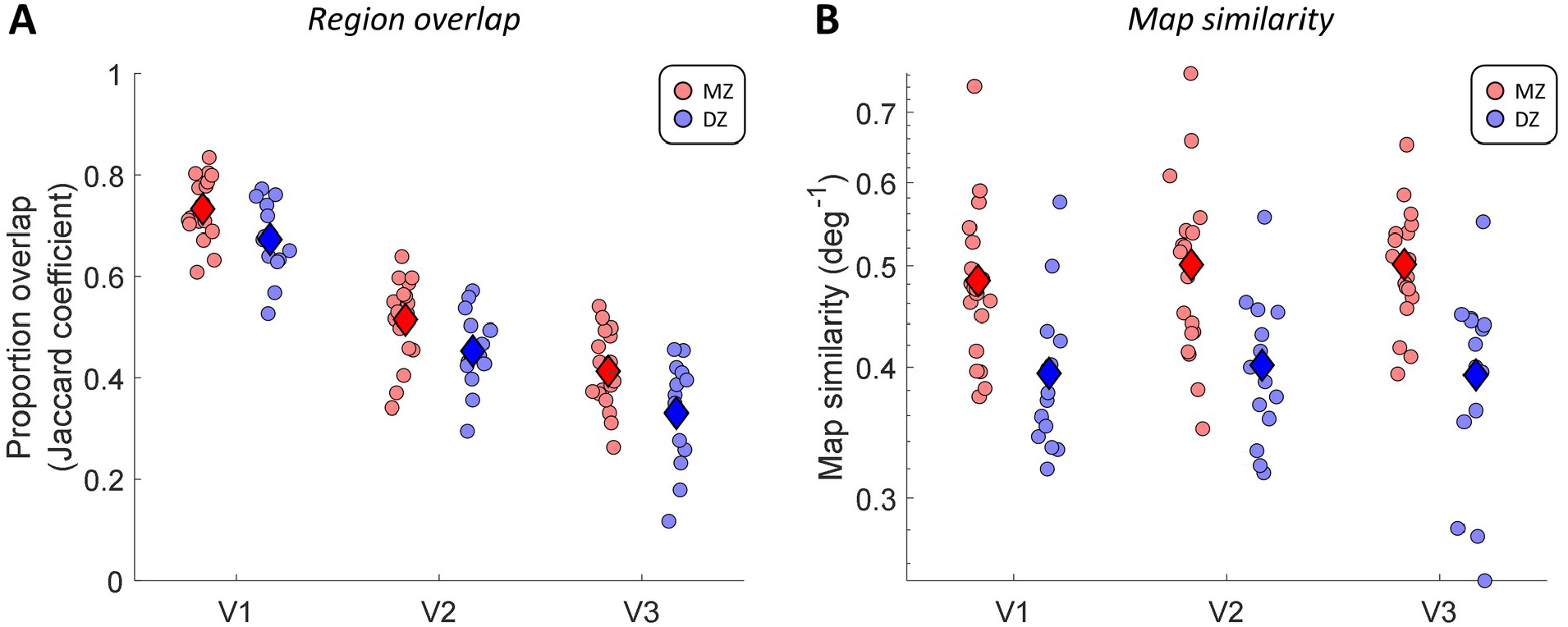
Proportion of overlap (A) and multivariate map similarity (B) for visual regions V1-V3 between twins in each pair. Each dot denotes the results for one twin pair in each cortical region. Diamonds indicate the group means. Map similarity scores were calculated as the inverse of the Euclidean distance in a multivariate space of pRF parameters (see 2.2.6). Note that map similarity scores are shown on logarithmic scale. Statistical inference was conducted using the logarithm of this score. Red: MZ twins. Blue: DZ twins.

We further quantified the significance of these overlap statistics using a permutation analysis. We shuffled the individual participants 1,000 times and thus compared each participant to a pseudo-randomly chosen participant instead of their twin. We conducted this analysis separately for each twin group. For each visual region and twin group, we then calculated the proportion of permutations in which the overlap between pseudo-random pairs was equal to or greater than the overlap measured in the twin pairs. This effectively determines the probability of observing the level of overlap we found between twins in unrelated participants. While the overlap was significantly above this level for MZ twins in all cortical regions (V1-V3 all p<0.001), for DZ the overlap was not significantly different in any region (V1: p=0.325; V2: p=0.232; V3: p=0.429).

### 3.2. Heritable functional architecture in early visual regions

Our findings offer the first evidence that the general topology of visual regions is more similar in MZ than DZ twins. However, manual delineation is susceptible to errors resulting from variable signal-to-noise ratios, missing data, errors in spatial alignment, or artifacts in the pRF analysis. More importantly, the above analyses only quantified the macroscopic overlap of visual regions.

We therefore examined the heritability of the functional architecture *within* each visual region, as revealed by population receptive field (pRF) analysis. For this we again used retinotopic maps that were spatially aligned to a common template brain. We calculated a map similarity score (inverse of mean Euclidean distance) that considers all three pRF parameters: x- and y-position, and pRF size. The results again showed that maps for MZ twins are substantially more similar than those in DZ twins (Figure 2B). We found maps were significantly more similar between MZ twins (F(1,93)=41.33, p<0.0001), but no significant effect of visual region (F(2,93)=0.18, p=0.8372) or any interaction between those factors (F(2,93)=0.23, p=0.7986).

To interpret these findings, it is important to ensure that these pRF measures are plausible. As well as visual inspection to ensure the retinotopic maps obtained from individual participants showed the expected structure in occipital cortex (as seen in Figure 1 and the Supplementary Information), we also quantified this formally in two ways. First, we plotted the relationship between mean pRF size and eccentricity, binned into 1 degree wide eccentricity bands, for each participant and visual region (Supplementary Figure S3). All participants showed a typical range of pRF sizes for each region that is consistent with those reported in the literature. Moreover, the group average for the two twin groups was very similar.

In addition, we calculated the mean visual field coverage for each twin group and visual region (Supplementary Figure S4). This measure effectively averages all the Gaussian pRF profiles from a given region in visual space, and thus quantifies how densely a given location in visual space is sampled by pRFs. Qualitatively, the coverage plots for MZ and DZ twins are very similar. Coverage in the periphery increases somewhat from V1 to V3, likely in accordance with increasing pRF size.

### 3.3. Heritability of individual pRF parameters

To unpack which pRF parameters drive the similarity of MZ maps, we quantified the degree of similarity in the spatial distribution of pRF parameters separately. This analysis also used maps that were spatially normalized to the common template. Specifically, we extracted the best-fitting polar angle and eccentricity preferences across vertices, as well as pRF sizes, separately for each visual region, and used a two-stage, random-effects analysis to estimate the average intra-class correlation for each twin group (see Methods for details). Finally, we calculated the heritability, h^2^, using Falconer’s formula (Falconer, 1965) and used a bootstrapping test to determine the significance of heritability for each pRF parameter and visual region.

For each pRF parameter, the first stage of the analysis computed the correlation between the parameter values in each twin pair within each visual region. The second stage then assesses heritability by testing whether the average similarity for MZ twins is greater than that for DZ twins. This analysis effectively treats each twin pair as a replication, meaning that the correlation for each twin pair is based on thousands of cortical surface vertices as observations. To illustrate the rationale of this approach, consider the analogy of comparing facial structure between twin pairs. For each twin pair, one could quantify facial similarity e.g. by taking a picture of each individual’s face and calculating heritability for each pixel in the image (see e.g. Tsagkrasoulis et al., 2017). This would be analogous to estimating heritability at each individual vertex in the cortical surface in our data – an approach that would require an infeasibly large sample for single-site fMRI study with twin pairs. An alternative approach would be to correlate intensity values across all pixels for a given pair of twins. In a second step, these correlations could then be compared between pairs of MZ and DZ twins to estimate the overall heritability of facial similarity. This would be analogous to our approach, where we correlate pRF parameter values across all vertices from a given visual region and then compare these correlations between twin groups.

By utilizing both within- and between-subject variations, we can therefore estimate heritability with a high degree of precision. To quantify this formally, we conducted a power analysis by simulating a range of plausible true heritability levels based on V3, the smallest visual region we analyzed. This showed that our analysis is well conditioned with 80% power to detect a heritability of approximately 3-4% (Supplementary Figure S2).

Our results suggest a degree of heritability for all three pRF properties – their preferred polar angle and eccentricity values, and their size. Figure 3A-C shows the average intra-class correlation for the three pRF properties across the three visual regions V1-V3. The degree of heritability can be appreciated by visual inspection of these plots. If a parameter were determined entirely genetically, the MZ correlation would be 1 and the DZ correlation 0.5, because DZ twins only share 50% of their genes on average. However, even a strongly heritable trait can usually be partly attributed to unique variance (such as environmental factors). In that case, the MZ correlation will fall below 1, and the DZ correlation would be half the MZ correlation. The solid lines in Figure 3A-C therefore have a slope of 0.5 and denote the correlations one would expect if the parameter is heritable. This is in contrast to the dashed identity line, which plots the 1:1 correspondence between MZ and DZ twins that would arise if these parameters were driven to the same extent across individuals by non-genetic factors. We find values that generally fall between these extremes, suggesting a significant effect of heritability on all pRF properties tested.

**Figure 3.**
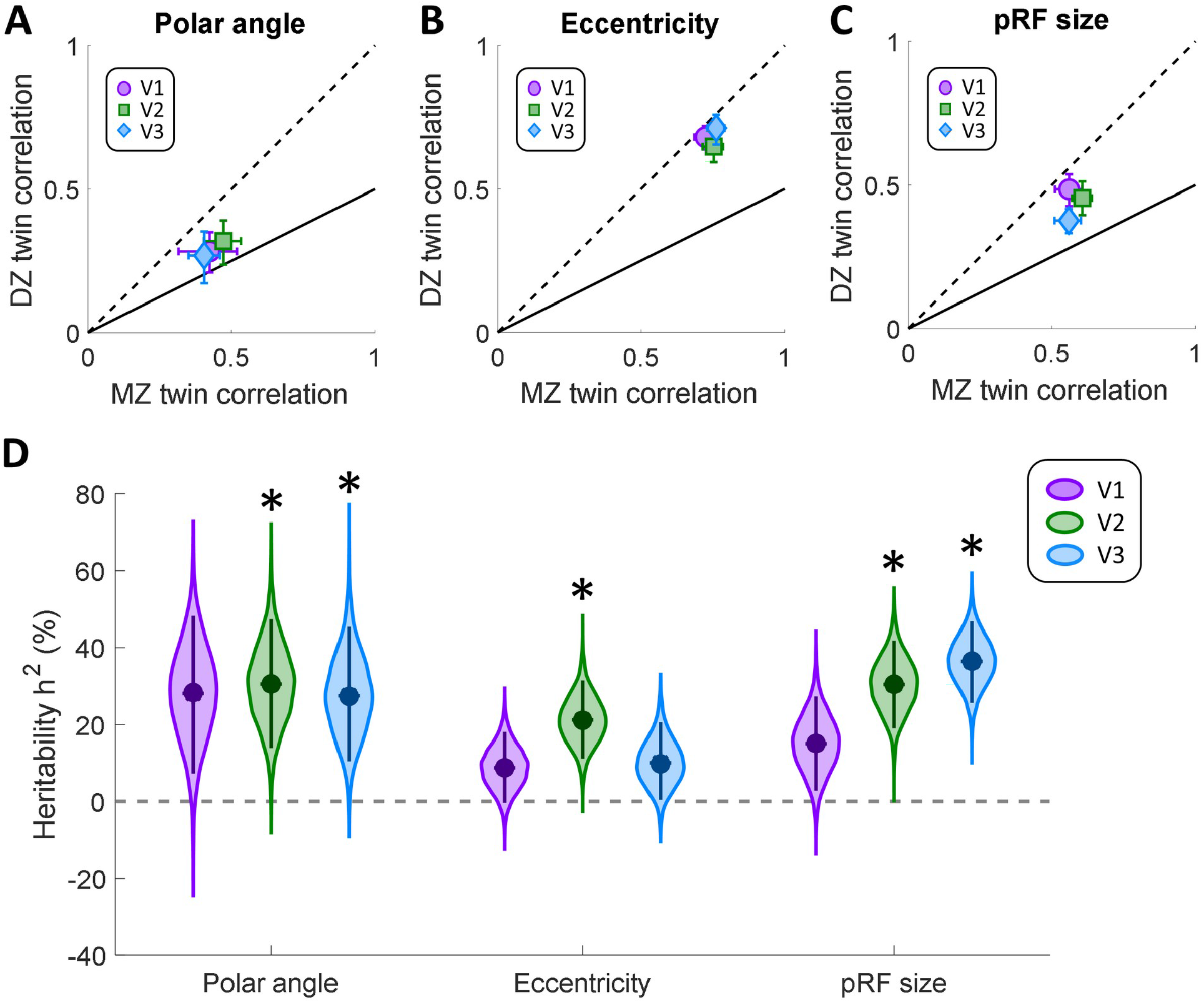
Intra-class circular correlations for polar angle (A), and Spearman correlations for eccentricity (B), and pRF size (C). MZ twin pair correlations are plotted against those for DZ twins. The solid line denotes the expected correlation if the variance was determined by genetic factors (see text for further details). The dashed line is the identity line. Error bars denote 95% confidence intervals derived through 10,000 bootstrap samples. D. Heritability for population receptive field parameters in each visual region. Data are shown for polar angle, eccentricity, and pRF size parameters, as derived from pRF analysis. Filled circles indicate the group means. The violin plot shows the bootstrap distribution for each pRF property and visual region, and the error bars denote 95% confidence intervals. Asterisks indicate significant differences at p<0.05, after Bonferroni correction for multiple comparisons.

Specifically, for polar angle (Figure 3A) and pRF size (Figure 3C), correlations were moderate (mean r range: 0.27-0.61), suggesting a high amount of unique variance in the members of each twin pair. Correlations were nonetheless stronger for MZ than DZ twins, indicative of some heritability. In contrast, correlations for eccentricity were generally strong (Figure 3B; mean r range: 0.65-0.76) but at similar levels for MZ and DZ twins, indicative of lower heritability.

We calculated these correlations after subtracting the average pattern across all participants, which allowed us to remove general trends such as the eccentricity gradient. Nevertheless, we further used a permutation analysis to examine if any of these general patterns were driving these results. We recalculated the correlations after shuffling the participants in each group 1,000 times to break up the twin pairs. Then, we determined the significance of these correlations by quantifying the proportion of resamples in which the mean intra-class correlation in shuffled pairs was equal to or greater than the mean correlation actually observed in the twin pairs. This measures the probability of these correlations occurring in unrelated participants. Interestingly, this analysis revealed that correlations for MZ twins were significantly higher than baseline for all pRF parameters and in all regions (all p≤0.005). In contrast, the correlations for DZ twins were not significantly greater than would be expected for unrelated participants (polar angle: all p≥0.105; eccentricity: all p≥0.120; pRF size: all p≥0.435).

Quantification of heritability using Falconer’s formula (Figure 3D) confirmed the above pattern. Polar angle preferences were significantly heritable (α_corrected_=0.0056) in V2 (h^2^=31%, p=0.0011) and V3 (h^2^=28%, p=0.0032), but not in V1 (h^2^=28%, p=0.0133). Heritability for eccentricity preferences was generally lower, and was significant only in V2 (h^2^=21%, p=0.0003), failing to reach significance in V1 (h^2^=9%, p=0.0582) and V3 (H^2^=10%, p=0.0421). Finally, pRF size was significantly heritable in V2 (h^2^=30%, p<0.0001) and V3 (h^2^=36%, p<0.0001), but not V1 (h^2^=15%, p=0.021).

We further analyzed the increase in heritability across the visual hierarchy by fitting a linear regression to each bootstrap iteration of heritability values for the three regions and determining the statistical significance of this change based on the bootstrapped slopes. This showed that the heritability for pRF size increased significantly from V1 to V3 (p=0.0044, Bonferroni corrected α=0.0167), while there was no such increase for polar angle (p=0.5277) or eccentricity (p=0.43).

When considering patterns of similarity between individuals, it is worth noting that BOLD signals from neighboring voxels, or vertices on the cortical surface, are not statistically independent (Logothetis, 2002). pRF estimates are more similar between neighboring than distal vertices, both due to the point spread function of the fMRI signal, and as a consequence of the gradient representation of the visual field in retinotopic maps. We therefore first subtracted the group average from individual maps. Nevertheless, a certain degree of correlation in pRF parameters is still expected between unrelated participants, which is borne out in our control analyses. However, spatial dependency cannot trivially account for the heritability results discussed thus far, because our results indicate that MZ twins display higher similarity in pRF parameters than DZ twins, or unrelated individuals. To ensure spatial similarity did not bias our results, we further conducted an analysis where instead of considering correlations between the complete retinotopic definition of each visual region, we sub-selected vertices ensuring a minimum separation of 8 mm between spatial points, to ensure non-adjacency. We then repeated the correlation analysis on this subset of vertices. Results were comparable to those from our main analysis (Supplementary Figure S5), although note that this analysis has of course only limited statistical power because of the substantially reduced number of data points.

Thus, our results demonstrate that visual cortical architecture is partially heritable – both in the topographic location of visual regions, and in the broad variations of pRF parameters, particularly polar angle and pRF size.

### 3.4. Heritability of cortical morphology

Spatial normalization from native brain space into a common template exploits cortical folding patterns to bring cortical surfaces into alignment. We should therefore expect brains to show a relatively high degree of similarity after normalization. It has been suggested that cortical morphology is linked to retinotopic map topology (Rajimehr and Tootell, 2009; Van Essen, 1997), and this is in fact a prerequisite for the creation of average and template maps (Benson et al., 2012; Sereno et al., 2013). How does similarity in cortical morphology impact the assessment of functional map heritability? To formally quantify the similarity of the structural measures in our twin pairs, we repeated the same analysis used for pRF parameters with anatomical parameters – namely, the cortical curvature and thickness estimates of each vertex derived by the FreeSurfer surface reconstruction algorithm.

Unsurprisingly, this analysis suggests a high degree of heritability for both parameters (Figure 4). The intra-class correlation plots show that the correlation coefficients fell below the identity line for all regions and both measures (Figure 4A-B). The correlations for V2 and V3 curvature were generally much weaker than for V1. This could be due to greater variability in the curvature of V2 and V3 given that these regions are smaller and comprise quadrant field maps, each of which fall approximately onto a single sulcal bank. In contrast, V1 is large and contains the calcarine sulcus. For thickness, V1 and V2 generally showed stronger correlations than V3.

**Figure 4.**
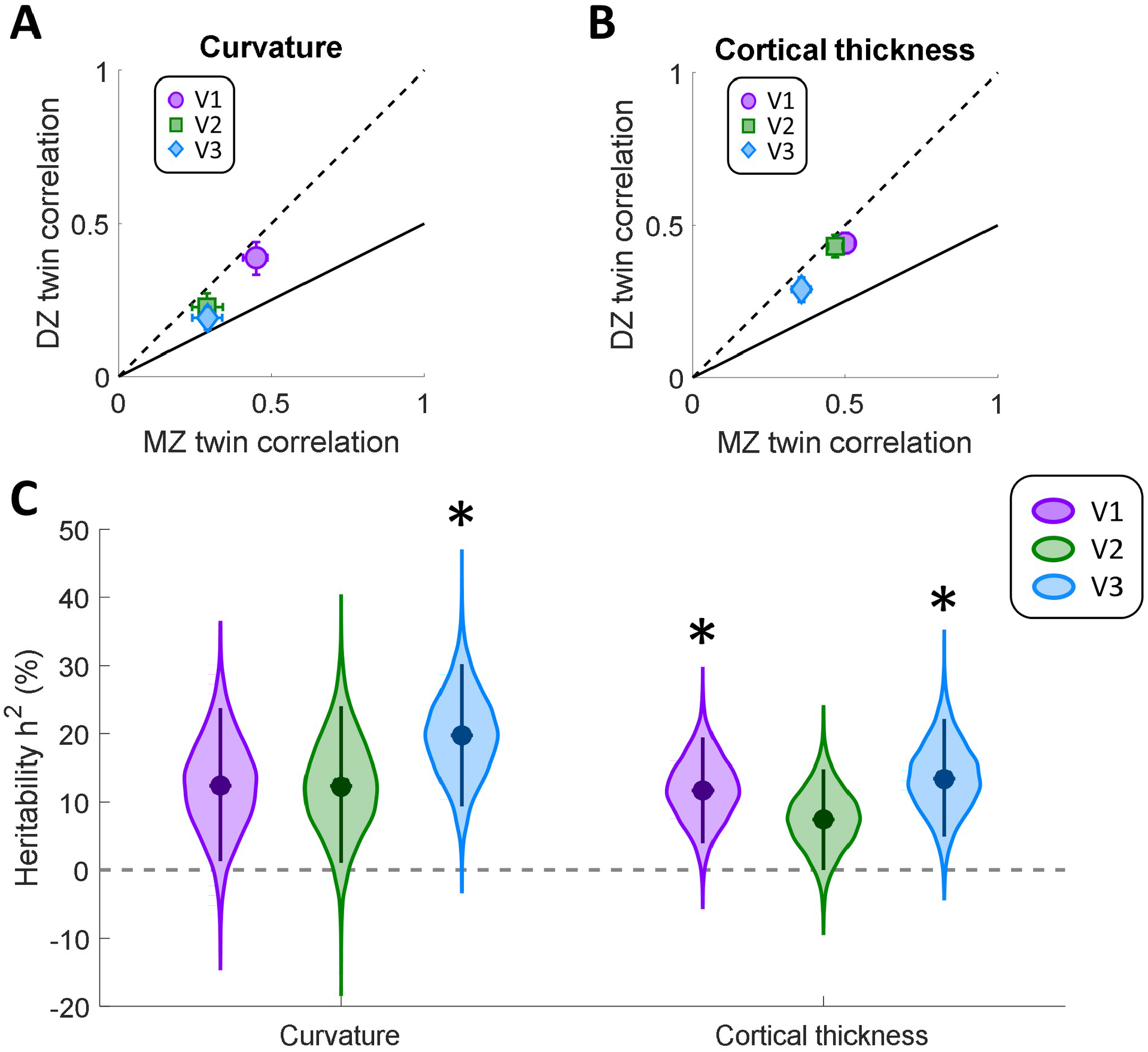
Intra-class Spearman correlations for cortical curvature (A), and thickness (B). MZ twin pair correlations are plotted against those for DZ twins. The solid line denotes the expected correlation if the variance was determined by genetic factors (see text for further details). The dashed line is the identity line. Error bars denote 95% confidence intervals derived through 10,000 bootstrap samples. C. Heritability for these anatomical parameters in each visual region. Data are shown for cortical curvature and thickness, as derived from surface reconstruction in FreeSurfer. Filled circles indicate the group means. The violin plot shows the bootstrap distribution for each pRF property and visual region, and the error bars denote 95% confidence intervals. Asterisks indicate significant differences at p<0.05, after Bonferroni correction for multiple comparisons.

Quantifying the heritability using the same statistical approach as for pRF parameters (Figure 4C), we found significant heritability (α_corrected_=0.0083) for cortical curvature only in V3 (h^2^=20%, p=0.0012), but not V1 (h^2^=12%, p=0.0345) or V2 (h^2^=12%, p=0.0379). For cortical thickness we found significant heritability in V1 (h^2^=12%, p=0.0068) and V3 (h^2^=14%, p=0.0047), but not V2 (h^2^=7%, p=0.0503).

We also conducted permutation tests to assess whether correlations for these cortical morphology parameters differed significantly from what can be expected from unrelated participants. This confirmed that intra-class correlations for MZ twins were highly significant for both cortical curvature and thickness in all three regions (all p<0.001). In contrast, for DZ twins correlations were not significantly greater than baseline (curvature: all p≥ 0.049, thickness: all p≥0.240). Taken together, our findings demonstrate that cortical morphology in early visual regions is partly heritable, in particular in V1 and V3.

### 3.5. Magnitude of spatial transformation

As described above, the greater structural similarity between MZ than DZ twins cannot be trivially explained by the effect of spatial normalization. If anything, this may artifactually increase the similarity of DZ brain structure given their greater dissimilarity to begin with. It could nonetheless be argued that because MZ brains are more similar, they also require a more similar spatial transformation to be warped into the common template. In order to test if residual differences in MZ are smaller because similar spatial transformation was applied to their maps, we quantified the average spatial transformation needed to align each brain to the template. Specifically, for each vertex in the sphere model (used for cortical alignment) we calculated the Euclidean distance from its native coordinates to the one used to align it with the template, and then averaged this for each visual region in each participant. Finally, we compared the amount of transformation in each region between twins by computing a correlation between them.

The results (Figure 5) show that the correlation between transformation statistics in MZ twins was very weak and not statistically significant (V1: r=-0.06, p=0.805, V2: r=-0.02, p=0.929, V3: r=0.05, p=0.833). If our results were to be driven by the similarity in the transformations applied to each twin in the pairs, we would expect to see greater correlations between these transformations in the MZ than the DZ twins. However, if anything, these transformations are slightly more correlated (albeit also non-significantly) in DZ twins (V1: r=0.19, p=0.524, V2: r=0.11, p=0.700, V3: r=0.01, p=0.983). We conclude that there is a low likelihood that our heritability estimates were driven by spatial normalization.

**Figure 5.**
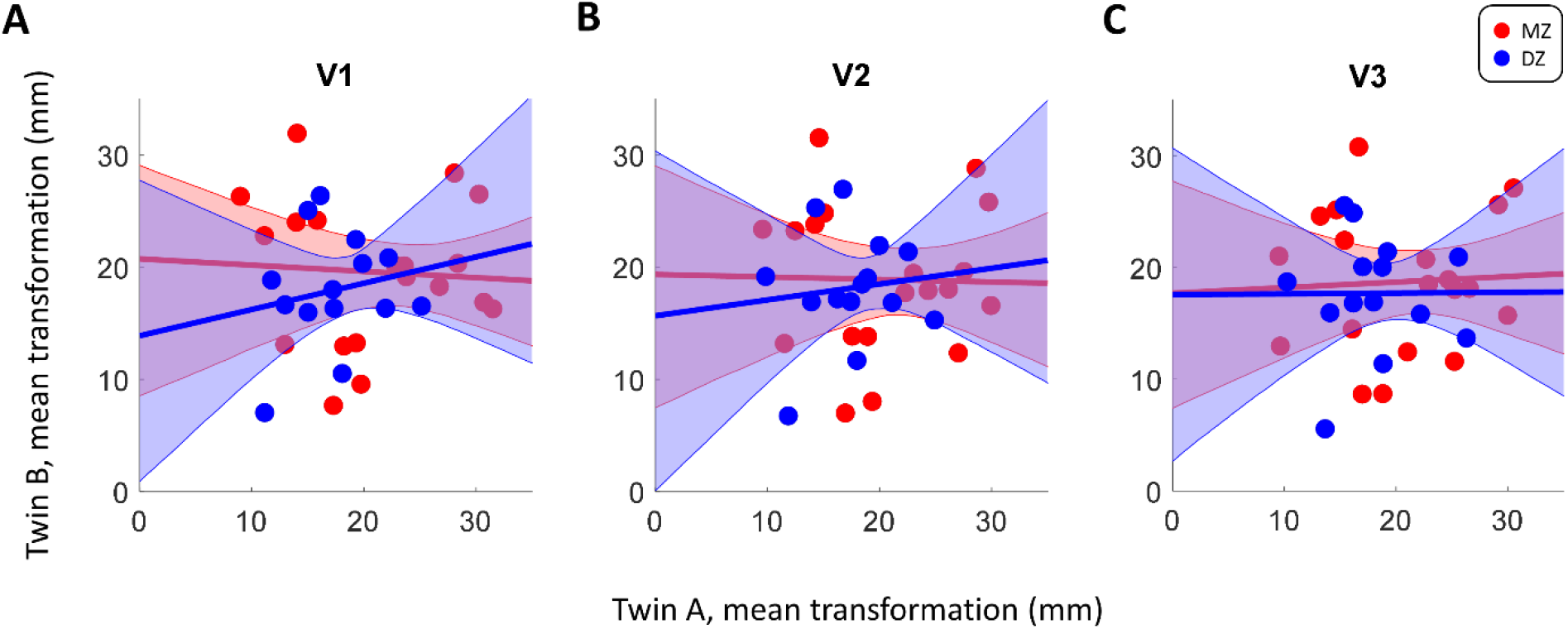
Comparison of the mean spatial transformation needed to align native brains to the template brain, separately for V1 (A), V2 (B), and V3 (C). Each dot denotes for a given twin pair the mean Euclidean distance between native vertices in a region and their location after cortical alignment. The solid lines denote the best fitting linear regression and the shaded regions shows the 95% confidence interval. Red: MZ. Blue: DZ.

## 4. Discussion

We examined whether the topology of early visual cortex has a genetic component by conducting fMRI testing on identical and non-identical twin pairs. Our findings suggest that both the cortical morphology and the functional architecture of early visual regions are at least partly driven by genetic components.

Our analysis revealed a moderate genetic component for the architecture of retinotopic maps, most notably for pRF sizes, a measure of the spatial selectivity of neural populations. The heritability factor for both pRF size and preferred polar angle ranged from 15-36% across regions of visual cortex. This suggests that at least two thirds of the variance in these measures was determined by environmental factors and random fluctuations. Interestingly, we observed strong correlations for pRF eccentricity preferences within twin pairs, irrespective of MZ or DZ status. This corresponds to a weak heritable component of 9-21% for eccentricity gradients in retinotopic organization, suggesting that a large proportion of the variance in eccentricity preferences may be due to other factors, including experience-dependent plasticity and/or constraints set by other processes in cortical development. Alternatively, our power to detect heritability of eccentricity may have been reduced due to the high overall correlations between maps. The MZ correlations observed here are close to the intersession reliability of eccentricity estimates for the same individual (van Dijk et al., 2016). Thus, if true MZ correlations between eccentricity values are higher, our method would not be sensitive to this and may have underestimated the heritability of eccentricity preferences as a result. Note, however, that the intersession reliability for polar angle is similarly strong (van Dijk et al., 2016), rendering our analysis sensitive to MZ-DZ differences because of the lower overall correlations in this parameter (Fig. 3A and C).

Given recent observations that eye movement patterns are heritable (Constantino et al., 2017; Kennedy et al., 2017), it is possible that eye movement behavior and retinotopic organization are linked. Indeed, the association between individual differences in eye-movement behaviors and a range of visual abilities (de Haas et al., 2019; Greenwood et al., 2017) suggests that differences in visual foraging behaviors, perhaps even in early postnatal life, could result in the formation of similar cortical representations of the visual field. Alternatively, the arrow of causality could point in the opposite direction: cortical idiosyncrasies in how the cortex encodes the visual field (Moutsiana et al., 2016) could determine where an observer directs their gaze. For instance, an observer may tend to foveate a given peripheral location with particularly poor resolution (large receptive fields) to bring it into clear view. Given that these idiosyncrasies are shared between MZ twins, this could result in more similar eye movements, as well as patterns of visual function across the visual field. This is an exciting possibility that could be investigated in future research.

Importantly however, the heritability of eye movements cannot trivially account for our present findings as a confounding factor during the experiment. Retinotopic mapping requires participants to maintain stable fixation on a central target, to ensure the accurate and systematic stimulation of different visual field locations. Nonetheless, eye movements could in principle influence our estimates of cortical selectivity via fixation stability – poorer fixation stability would result in a lower signal-to-noise ratio and poorer quality maps. Thus, two participants with more stable fixation would likely have more similar maps. We are not aware of evidence suggesting that fixation stability is heritable (as opposed to patterns of free viewing in complex visual scenes). Critically, to account for our findings the fixation stability of MZ twins would not have to be more similar to each other, but better than that of DZ twins in general. Furthermore, our results cannot be explained by more similar salience biases between MZ than DZ twins (e.g. for faces, Kennedy et al., 2017). The order of carrier images during our mapping runs were randomized between participants, meaning that variations in image salience could not have resulted in systematic mapping biases.

In all our analyses, we used retinotopic maps that were spatially aligned to a common template brain. To compare them directly, it is necessary to bring the data from individual participants into register. Importantly, this spatial distortion could not have artifactually produced our results. Based on previous research (Anderson et al., 2021; Chen et al., 2011; Jansen et al., 2015), we expect MZ brains to have more similar architecture than DZ brains, a finding that we also replicate. However, it does not follow that retinotopic maps should therefore become more similar after normalization. This notion rests crucially on the assumption that retinotopic maps are directly related to cortical morphology – that is, the hypothesis we set out to test. Rather, the opposite is the case: the similarity of DZ maps should artifactually increase due to spatial normalization. This makes our analysis more conservative. Finally, the amount of warping required to align individual brains to the template brain was uncorrelated between participants in either group, suggesting that MZ twins did not undergo a more similar amount of transformation, at least not in early visual cortex.

It is indeed likely that there are consistent patterns in retinotopic maps across all participants irrespective of zygosity or familial relationship. To control for this confound, we subtracted the grand average map across the whole sample from each individual map before conducting the correlation analysis on the residuals. However, this procedure only corrected for part of the general trend. It is possible that similar visual experiences during development, such as the exposure to urban versus rural settings, could result in plasticity that shapes retinotopic map organization. However, our permutation analysis using shuffled twin pairings suggested that the distributions of all pRF parameters were much more similar between MZ twins than would be expected between unrelated participants. This was not the case for DZ twins. Therefore, we surmise that variations in the visual environment of individuals in our sample likely only played a small role in shaping the structure of their retinotopic maps.

Our findings are noteworthy in the context of previously reported links between perception and properties of retinotopic cortex. Several studies have shown that activation patterns in V1 reflect the apparent size of stimuli (specifically, the apparent stimulus eccentricity) rather than the veridical size of the retinal image (Fang et al., 2008; Murray et al., 2006; Pooresmaeili et al., 2013; Sperandio et al., 2012). Further, the macroscopic surface area of V1 correlates with variations in perceived size from phenomena like the Ebbinghaus illusion (Bergmann et al., 2015, 2014; Schwarzkopf et al., 2011; Schwarzkopf and Rees, 2013; Verghese et al., 2014). The rate of alternations in binocular rivalry is also heritable (Miller et al., 2010). Interestingly, V1 surface area further predicts the temporal dynamics of travelling waves in binocular rivalry (Genç et al., 2014). Given that the macroscopic surface area of visual regions correlates with the spatial selectivity (Duncan and Boynton, 2003; Harvey and Dumoulin, 2011; Song et al., 2015, 2013), one would expect a link between these properties and the perceived alternations in rivalry. Altogether, these findings suggest a functional role for V1 and its retinotopic organization in a range of perceptual and cognitive processes. Our finding of heritable retinotopic organization therefore suggests that there should be a heritable component for these aspects of individual perception.

Our findings are consistent with recent demonstrations that the topology of functional connectivity networks is heritable, including in early sensory cortex (Anderson et al., 2021). The peak frequency of visually-induced gamma oscillations in visual cortex is also strongly heritable (van Pelt et al., 2012). We and others have reported links between gamma oscillation frequency, occipital levels of the neurotransmitter *gamma*-aminobutyric acid (Muthukumaraswamy et al., 2009), and the surface area of early visual cortex (Bergmann et al., 2016; Gregory et al., 2016; Schwarzkopf et al., 2012; but see also Cousijn et al., 2014). As a potential explanation for these links, we posited that gamma oscillation frequency depends on the cortical microarchitecture, and thus on the spatial selectivity of neuronal populations (Pinotsis et al., 2013; Schwarzkopf et al., 2012). Our present findings may constitute another missing link between these diverse aspects of visual processing: if cortical organization gives rise to gamma oscillations, then heritability in visual cortex architecture could also explain why these aspects are heritable.

The differentiation of cortex into specialized brain regions during development is driven both by intrinsic genetic mechanisms, such as signaling molecules, and extrinsic topographically organized afferents (O’Leary, 1989; O’Leary et al., 2007; Rakic, 1988; Rakic et al., 2009). In children as young as 6-7 years, both the organization and functional properties of visual cortex are already similar to adults (Conner et al., 2004; Dekker et al., 2019). The development of the Stria of Gennari is preserved in congenital blindness (Trampel et al., 2011), further supporting a strong genetic determination of the arealization in V1, at least. Nevertheless, there is great experience-dependent plasticity in the development of functional response properties of visual cortex (Hubel and Wiesel, 1965; Wiesel and Hubel, 1965a, 1965b). Early blindness causes structural changes in visual brain regions (Touj et al., 2020), possibly due to functional reorganization to serve other functions (Burton, 2003). Spatial selectivity of visual cortex, as measured by pRF size, is also abnormal in individuals with amblyopia (Clavagnier et al., 2015). Thus, it seems likely that the organization of retinotopic maps depends on both genetically determined processes and experience-dependent fine-tuning of neuronal selectivity.

Previous research has also suggested a close correspondence between cortical curvature and the structure of retinotopic maps, such that the vertical meridian tends to map onto the gyral lips and the horizontal meridian maps onto the sulcal depth (Rajimehr and Tootell, 2009). In fact, this correspondence allows researchers to spatially normalize brains and average maps as we did here, and even to leverage probabilistic atlases for identifying retinotopic regions without collecting functional data (Benson et al., 2012). The finding of heritable networks of functional connectivity between brain regions (Anderson et al., 2021) again suggests a role of genetics in determining cortical architecture. It is possible that the constraints placed on intra-cortical connections due to retinotopic map organization could determine cortical folding (Van Essen, 1997). This hints at the interesting possibility that the genetic component in retinotopic map development may also drive cortical folding, though this hypothesis will need to be tested explicitly in future research.

In conclusion, our study is the first to investigate the heritability of the functional organization of visual regions in the human brain. Future studies must seek to understand the implications of these aspects for visual processing and perceptual function. Previous research has demonstrated that the way each of us perceives the visual world is at least partly determined by genetics. Our findings hint at an exciting possibility that this is because the spatial architecture of our visual cortex is inherited.

## 5. Acknowledgments

This research was supported by European Research Council Starting Grants 310829-WMOSPOTWU (DSS) and 852885-INDIVISUAL (BdH), as well as the Deutsche Forschungsgemeinschaft grant 222641018-SFB/TRR 135 TP A8 (BdH). IA was supported by the Medical Research Council (MR/K014382/1) and by core funding from the Wellcome Trust (203139/Z/16/Z), and JAG by the Medical Research Council (MR/K024817/1).

## 6. Author contributions

IA: data analysis, consultation, revised manuscript

NF: conception, study organization, data collection, initial analysis and draft manuscript

SE: data collection, study organization, initial analysis, revised manuscript BdH: consultation, data analysis, revised manuscript

JAG: consultation, data analysis, revised manuscript

DSS: conception, data analysis, data curation, consultation, revised manuscript

## 7. Data availability

Normalized retinotopic mapping data and analysis code available at: https://doi.org/10.17605/OSF.IO/Q8DRF

## Appendix: Supplementary Information

**Figure S1.**
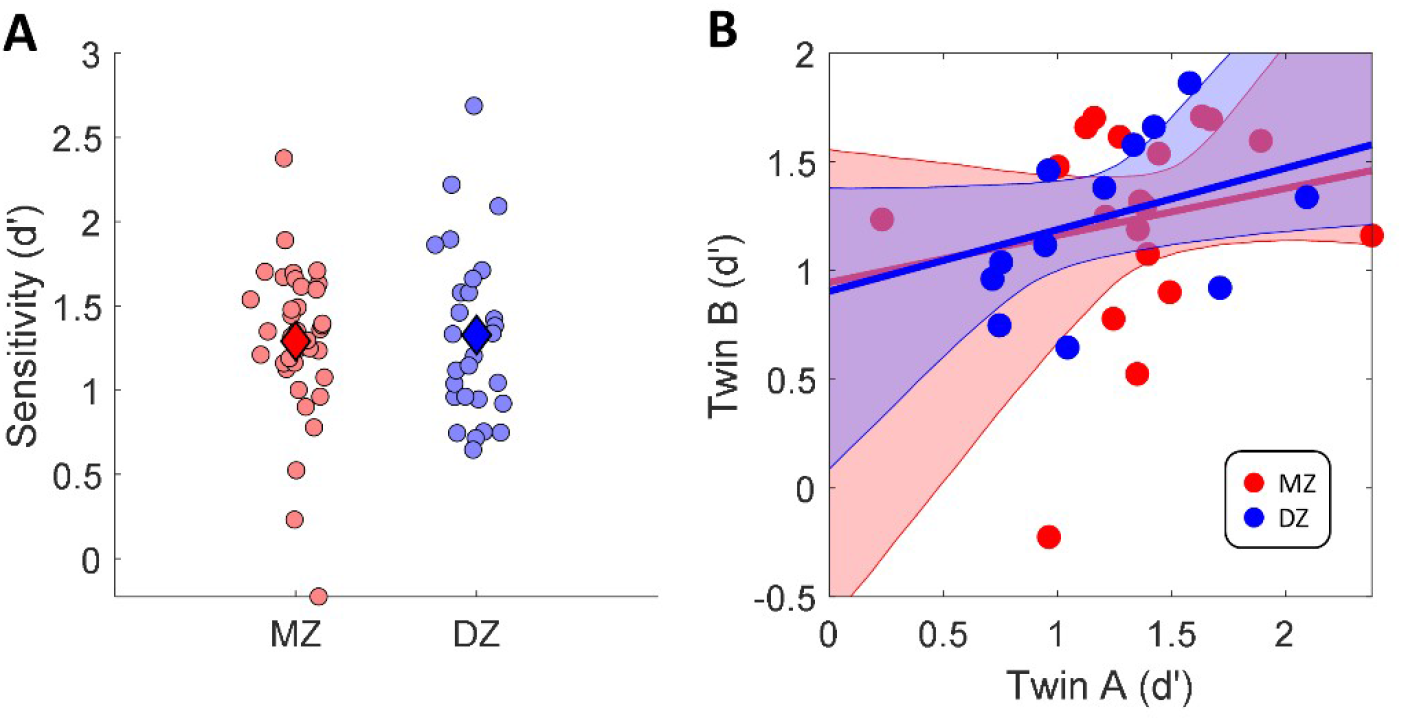
Behavioral results. A. Sensitivity (d’) on the behavioral task in the scanner for MZ and DZ twin groups. Each dot denotes the results for one twin pair in each cortical region. Diamonds indicate the group means. B. Comparing behavioral sensitivity (d’) between twins in each pair. Each dot denotes a given twin pair. The solid lines denote the best fitting linear regression and the shaded regions shows the 95% confidence interval. Red: MZ twins. Blue: DZ twins.

**Figure S2.**
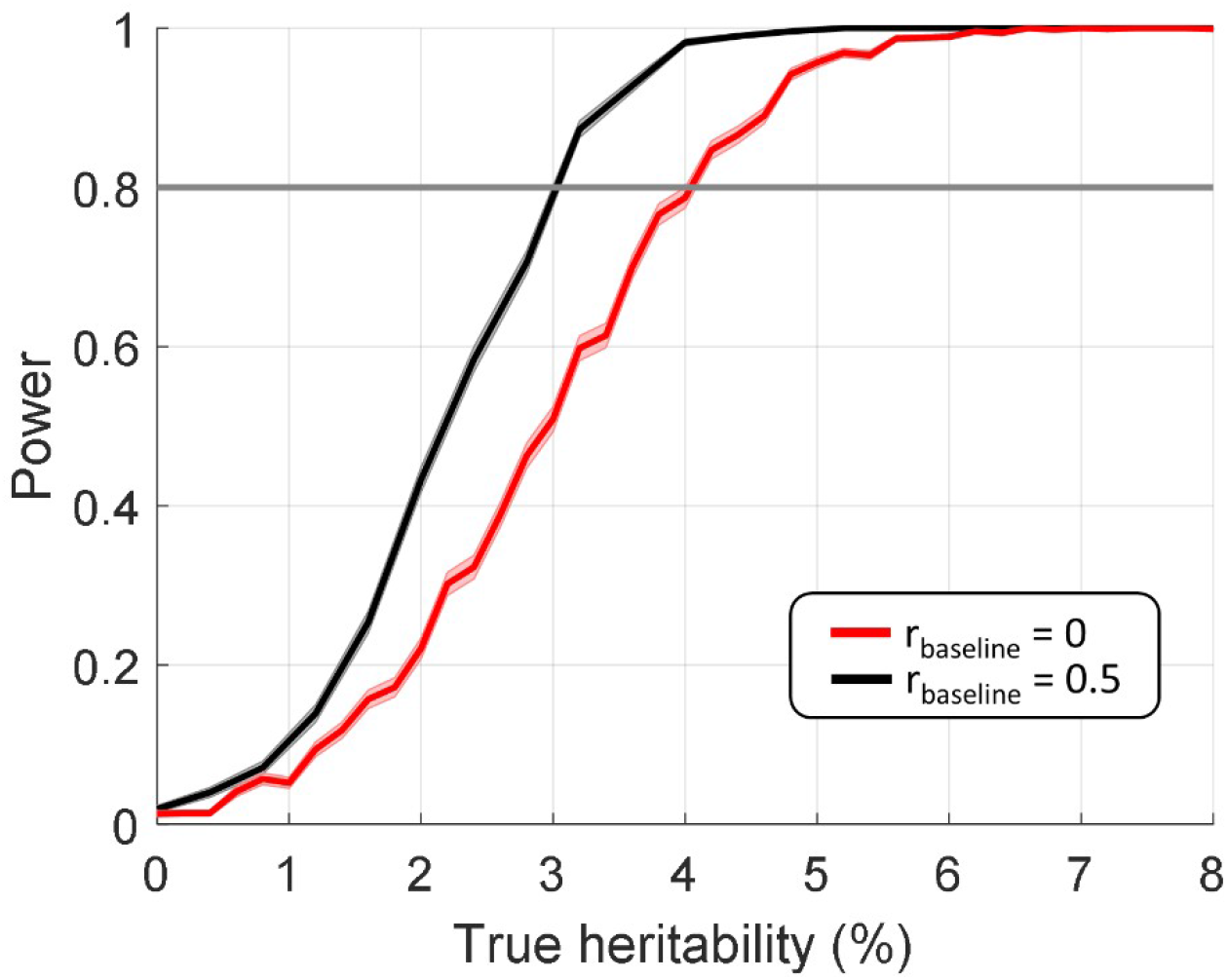
Power analysis for pRF parameter correlations. We simulated data with the same number of spatial points as the smallest visual region we report, area V3 (3409 vertices), and the same sample sizes for the MZ (n=19) and DZ (n=14) groups as in our study. We modulated the difference in correlation between the MZ and DZ groups to simulate a range of possible heritability values. We repeated this simulation 1,000 times. For each simulated heritability value, we then quantified the number of times that our bootstrapped heritability analysis (see Materials and Methods) detected a significant effect (p<0.05, Bonferroni corrected for 9 comparisons). The plot shows the detection rate (power) against true heritability. This suggests our analysis achieved 80% power to detect a true heritability of approximately 3-4%, depending on whether we assume a mid-range baseline intra-class correlation of r=0.5 or the minimal DZ correlation of r=0.

**Figure S3.**
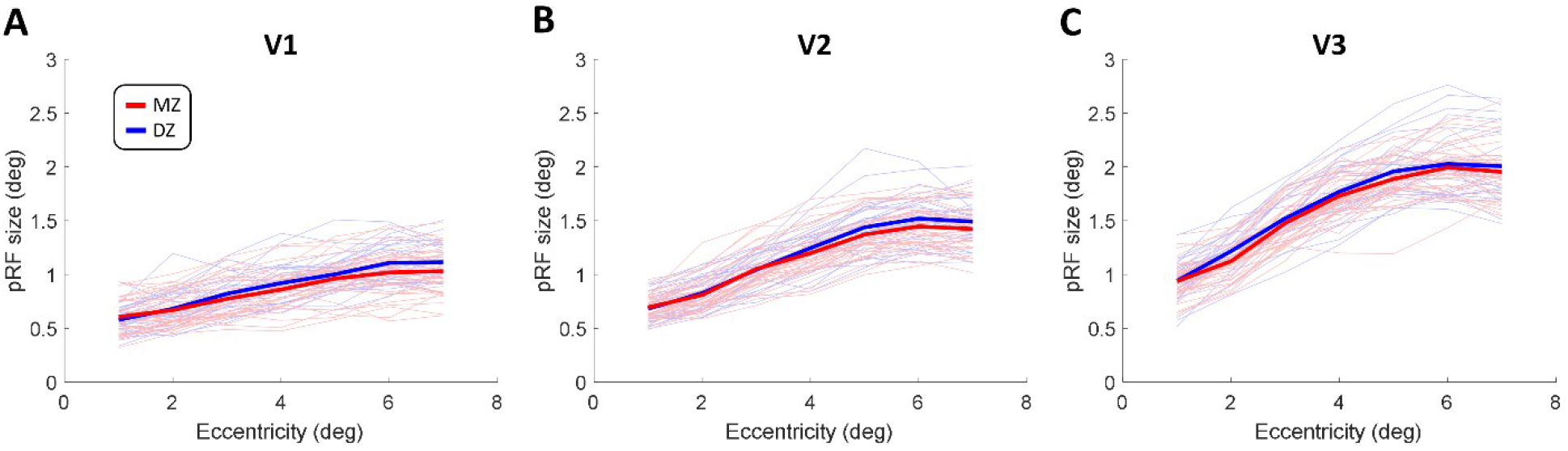
pRF size relative to eccentricity. pRF size data for each participant from V1 (A), V2 (B), or V3 (C) were binned into 1 degree wide eccentricity bins and averaged. Bin averages are plotted against eccentricity. The faint curves show individual participants. The thicker lines denote the mean of each twin group. Red: MZ, Blue: DZ.

**Figure S4.**
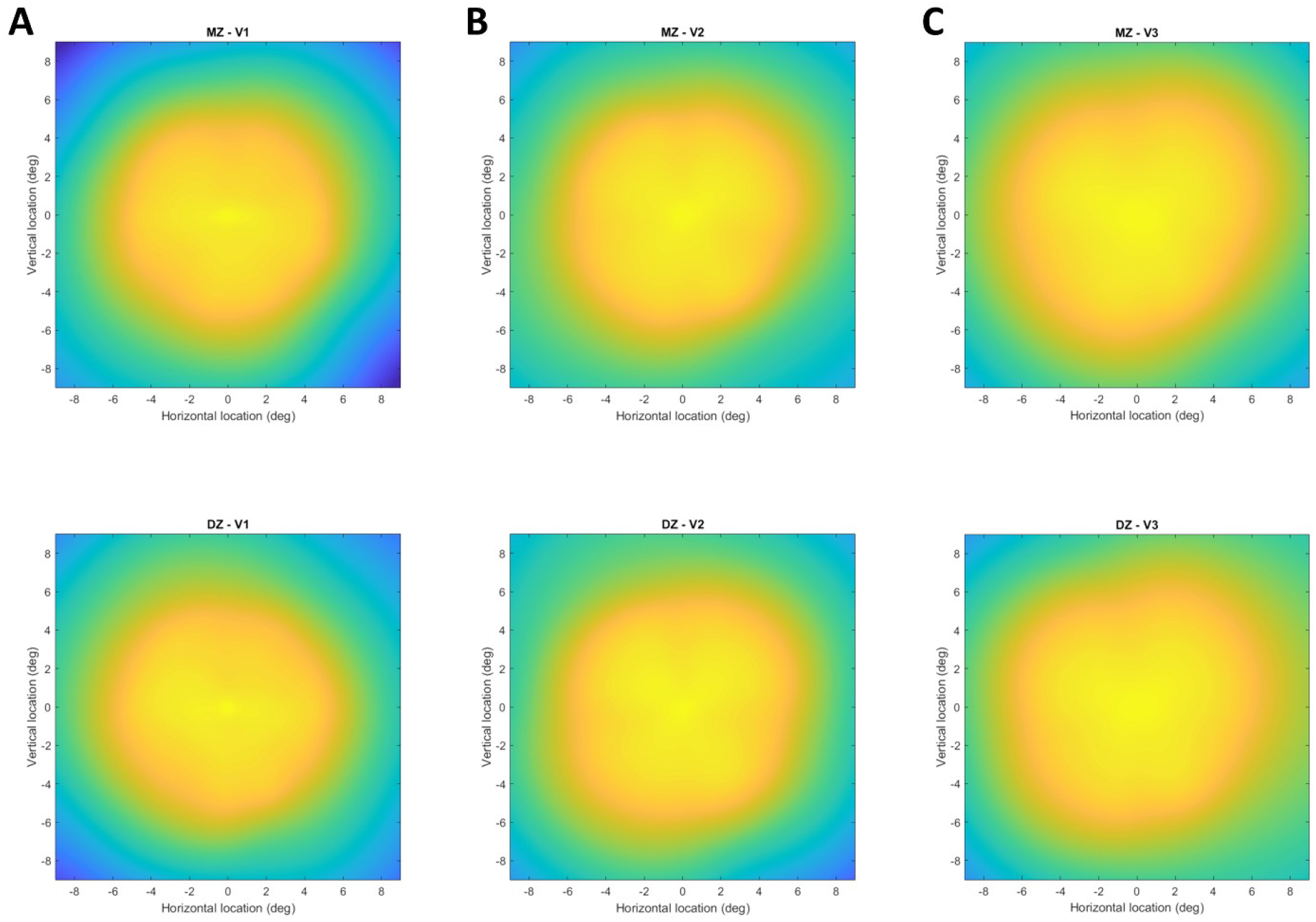
Visual field coverage plots for V1 (A), V2 (B), and V3 (C). Visual field coverage reflect the average of all pRF profiles within a region, thresholded with R^2^>0.1 and only including pRFs whose centers fell within 9 degrees eccentricity. Plots were further averaged across all participants in each group. The color of each pixel denotes how densely each visual field location was sampled by the pRFs in the region, ranging from blue (low coverage) over green to yellow (high coverage). Top row: MZ twins. Bottom row: DZ twins.

**Figure S5.**
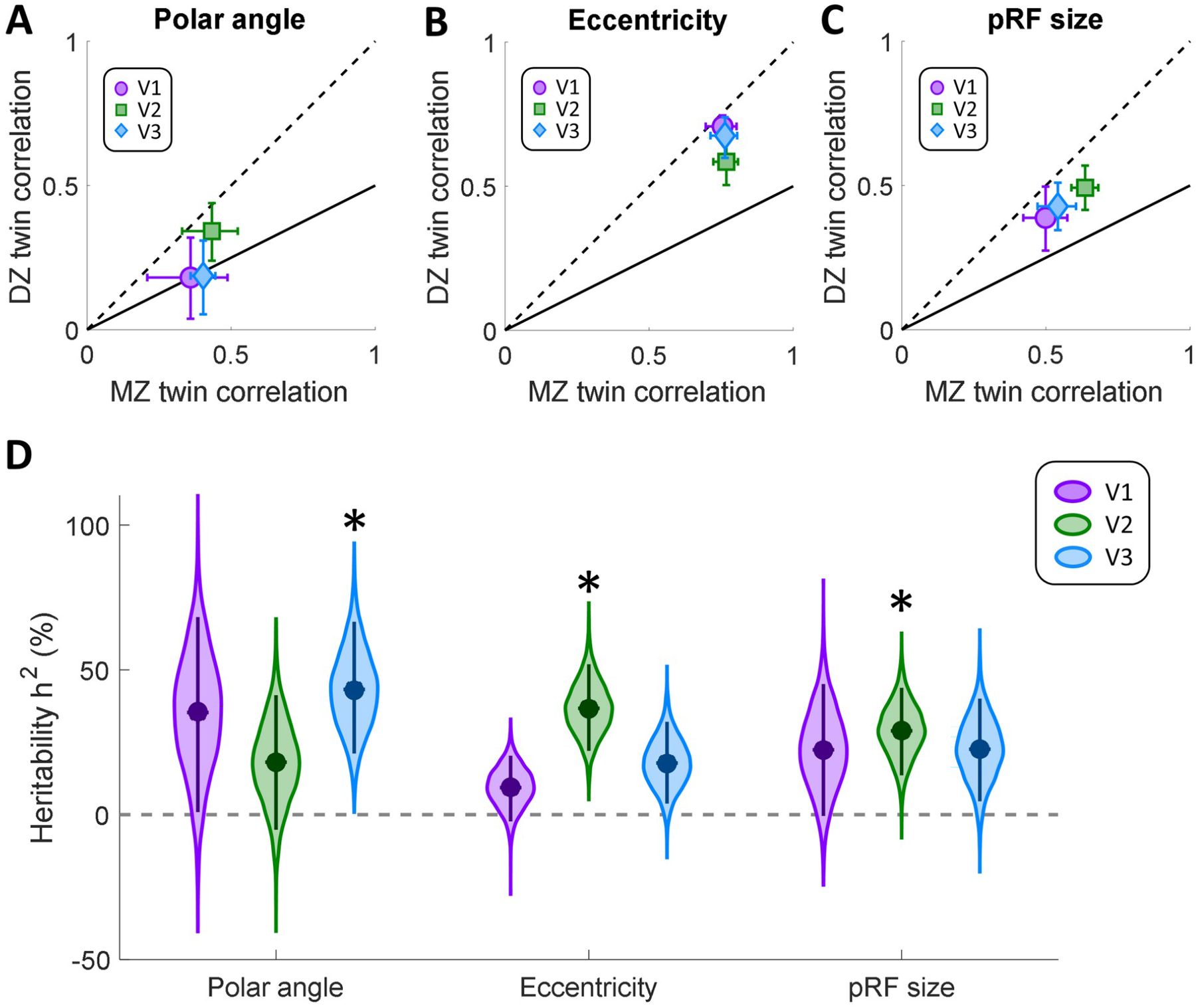
Control analysis of intra-class circular correlations for polar angle (A), and Spearman correlations for eccentricity (B), and pRF size (C). In order to provide an estimate that is less biased by the spatial dependence between neighboring vertices, we selected vertices with a minimal spatial separation of 8 mm between spatial sample points. MZ twin pair correlations are plotted against those for DZ twins. The solid line denotes the expected correlation if the variance was determined by genetic factors (see text for further details). The dashed line is the identity line. Error bars denote 95% confidence intervals derived through 10,000 bootstrap samples. D. Heritability for population receptive field parameters in each visual region. Data are shown for polar angle, eccentricity, and pRF size parameters, as derived from pRF analysis. Filled circles indicate the group means. The violin plot shows the bootstrap distribution for each pRF property and visual region, and the error bars denote 95% confidence intervals. Asterisks indicate significant differences at p<0.05, after Bonferroni correction for multiple comparisons.

## Notes

https://doi.org/10.17605/OSF.IO/Q8DRF

### Competing Interest Statement

The authors have declared no competing interest.

### Summary of Updates

How much of the functional organization of our visual system is inherited? Here we tested the heritability of retinotopic maps in human visual cortex using functional magnetic resonance imaging. We demonstrate that retinotopic organization shows a closer correspondence in monozygotic (MZ) compared to dizygotic (DZ) twin pairs, suggesting a partial genetic determination. Using population receptive field (pRF) analysis to examine the preferred spatial location and selectivity of these neuronal populations, we further demonstrate that across cortical regions V1-V3, map architecture was more similar in MZ than DZ twins. The heritability of spatial selectivity, as quantified by pRF size, increased across the visual hierarchy. Our findings are consistent with heritability in both the arrangement of visual regions and spatial tuning properties of visual cortex. This could constitute a neural substrate for variations in a range of perceptual effects, which themselves have been found to be at least partially genetically determined.

